# Cryo-EM structure of α-Synuclein Fibrils Harboring the Dementia with Lewy Bodies–Associated E83Q Mutation

**DOI:** 10.64898/2025.12.23.696295

**Authors:** Notash Shafiei, Inayatullah Mohammed, Senthil T. Kumar, Anne-Laure Mahul-Mellier, Babatunde Ekundayo, Daniel Stähli, Amanda J. Lewis, Henning Stahlberg

**Affiliations:** Laboratory of Biological Electron Microscopy, Institute of Physics, School of Basic Sciences, École Polytechnique Fédérale de Lausanne, Rt. de la Sorge, 1015 Lausanne, Switzerland; Department of Fundamental Microbiology, Faculty of Biology and Medicine, University of Lausanne, Rt. de la Sorge, 1015 Lausanne, Switzerland; Laboratory of Molecular and Chemical Biology of Neurodegeneration (LMNN), School of Life Sciences, Brain Mind Institute, Ecole Polytechnique Fédérale de Lausanne (EPFL), CH-1015 Lausanne, Switzerland; National Centre for Cell Science, Pune, India

## Abstract

Aggregation of α-synuclein (α-Syn) into amyloid fibrils underlies the pathology of synucleinopathies. Rare familial mutations can modulate α-Syn aggregation and may give rise to distinct fibril conformations linked to disease heterogeneity. The E83Q mutation, identified in a Dementia with Lewy Body disease (DLB) patient with atypical Lewy body distribution, has been shown to accelerate α-Syn aggregation and enhance neuronal seeding, yet its structural basis remains unclear. Here, we determined the cryo-electron microscopy structure at 3.4 Å resolution of recombinant full-length human α-Syn fibrils harboring the E83Q mutation. The fibrils adopt a double-protofilament architecture with a conserved Greek-key–like core spanning residues 36–99, stabilized by intra- and inter-filament salt bridges. Despite this conserved fold, the E83Q mutation induces a distinct local rearrangement within the hydrophobic region: substitution of Glu83 with Gln reorients residue 83 inward, abolishing solvent exposure and altering interactions with the N-terminal region. This structural shift brings the hydrophobic region closer to the N-terminus, shielding residues implicated in post-translational modification and ligand binding. Our findings reveal how a single charge-neutralizing mutation reshapes α-Syn fibril architecture, providing a structural framework to understand the enhanced aggregation and pathogenic properties associated with the E83Q variant.

## Introduction

The protein alpha-synuclein (α-Syn) has been found at high concentration in cellular inclusions such as Lewy bodies (LB) and Lewy neurites (LN). These are hallmarks of synucleinopathy diseases, which include Parkinson’s disease (PD), dementia with Lewy bodies (DLB) and multiple system atrophy (MSA)^1,2^. In its native form, α-Syn interacts with various cellular components, but under pathological conditions it forms aggregates of misfolded protein in the brains of patients with these diseases. α-Syn has been shown to aggregate into amyloid fibrils, which is thought to be a crucial molecular process in the pathogenesis of Parkinson’s disease and related α-Synucleinopathies^3–6^. Different polymorphs of α-Syn fibrils may contribute to the clinical heterogeneity of synucleinopathies^7,8^. Therefore, understanding the structure of α-Syn fibrils is important for the development of precision therapies and biomarkers.

While most cases of Parkinson’s disease occur sporadically and are not associated with a known genetic mutation, there are familial forms of the disease that can be linked to genetic factors. Known familial α-Syn mutations include A30G^9^, E46K^10^, H50Q^11^, A53E^12^, A53T^13^, and A53V^14^. These mutations provide pathological molecular perturbations that have been shown to alter α-Syn aggregation kinetics, fibril morphology, and toxicity. Despite extensive biochemical and cellular characterization, how individual disease-linked mutations reshape the three-dimensional architecture of α-Syn fibrils remains poorly defined.

Recently, Kapasi and colleagues reported a unique α-Syn mutation, substituting the negatively charged glutamic acid at position 83 with the neutral glutamine (E83Q), which was found in a patient diagnosed with DLB. Histologically, the brain of this patient had widespread LB pathology in the hippocampus and cortical regions, with sparse pathology in substantia nigra (SN), distinguishing it from other known genetic and sporadic DLB cases^15^. This atypical regional distribution suggests that the E83Q mutation may confer distinct pathogenic properties to α-Syn aggregates.

Later, Kumar et al. investigated the biochemical properties and toxicity of fibrils formed from this α-Syn mutant in neuronal cell cultures^16^. They demonstrated that the E83Q mutation accelerates α-Syn aggregation in vitro, yielding fibrils with unique conformational and morphological features and with increased neuronal seeding capacities compared to the wild-type (WT) protein. Notably, E83Q fibrils induced the formation of structurally and morphologically heterogeneous Lewy body–like inclusions in neurons, closely recapitulating the diversity of α-syn pathology observed in human synucleinopathies. These findings suggest that the E83Q mutation alters fundamental properties of α-Syn assemblies that govern strain behavior and pathogenic potential. However, the molecular and structural determinants underlying these altered fibril properties remained unresolved, motivating direct high-resolution structural analysis of E83Q fibrils.

Here, we used cryo-electron microscopy (cryo-EM) to investigate the effects of the E83Q mutation on the structure of recombinant full-length α-Syn fibrils. This mutation is in the hydrophobic region of α-Syn, formed by residues 61-95 and formerly called non-amyloid-β component (NAC) region^15^, which has been shown to be crucial for the fibrillization process. The computed map at a resolution of 3.4 Å shows clear side-chain densities for residues 36–99. The protofilament interface is stabilized by a salt bridge between E57 and K58, and an additional intra-filament salt bridge is present between K96 and D98 in the C-terminus. While the Greek-key-like core was preserved, E83Q exhibited structural differences compared to WT and other α-Syn fibril polymorphs. The substitution of GLU83 with GLN causes the side chain to orient inward, eliminating its solvent accessibility. This inward shift disrupts long-range hydrophilic interactions and repositions the hydrophobic region closer to the N-terminus, a feature not observed in WT or other mutant fibrils. These structural changes provide a plausible molecular basis for the distinct aggregation and seeding properties associated with the E83Q mutation.

Our findings provide a structural framework for understanding how single-point mutations within the NAC region remodel the α-synuclein fibril core to produce distinct architectures, offering a molecular basis for strain diversity and divergent neuropathological outcomes in synucleinopathies.

## Results

### Preparation of E83Q α-Syn fibrils

Recombinant full-length human α-Syn harboring the E83Q mutation was expressed and purified as previously described^17^. The purity and identity of the monomeric protein was confirmed by SDS-PAGE, UPLC and mass spectrometry analysis, which showed a single band and a molecular mass consistent with full-length E83Q α-Syn (**Supplementary Fig. 1A-C**). Negative stain TEM was used to screen fibrils formed under different conditions like temperatures, shaking or quiescent conditions (**Fig. 1A**). The fibrils assembled in 1X phosphate buffered saline (PBS) at pH 7.4 under continuous shaking at 900 rpm for 15 hours, showed a high proportion of twisted fibrils with a well-resolved crossover structure. These were selected for cryo-EM experiments (**Fig. 1B**).

**Figure 1:**
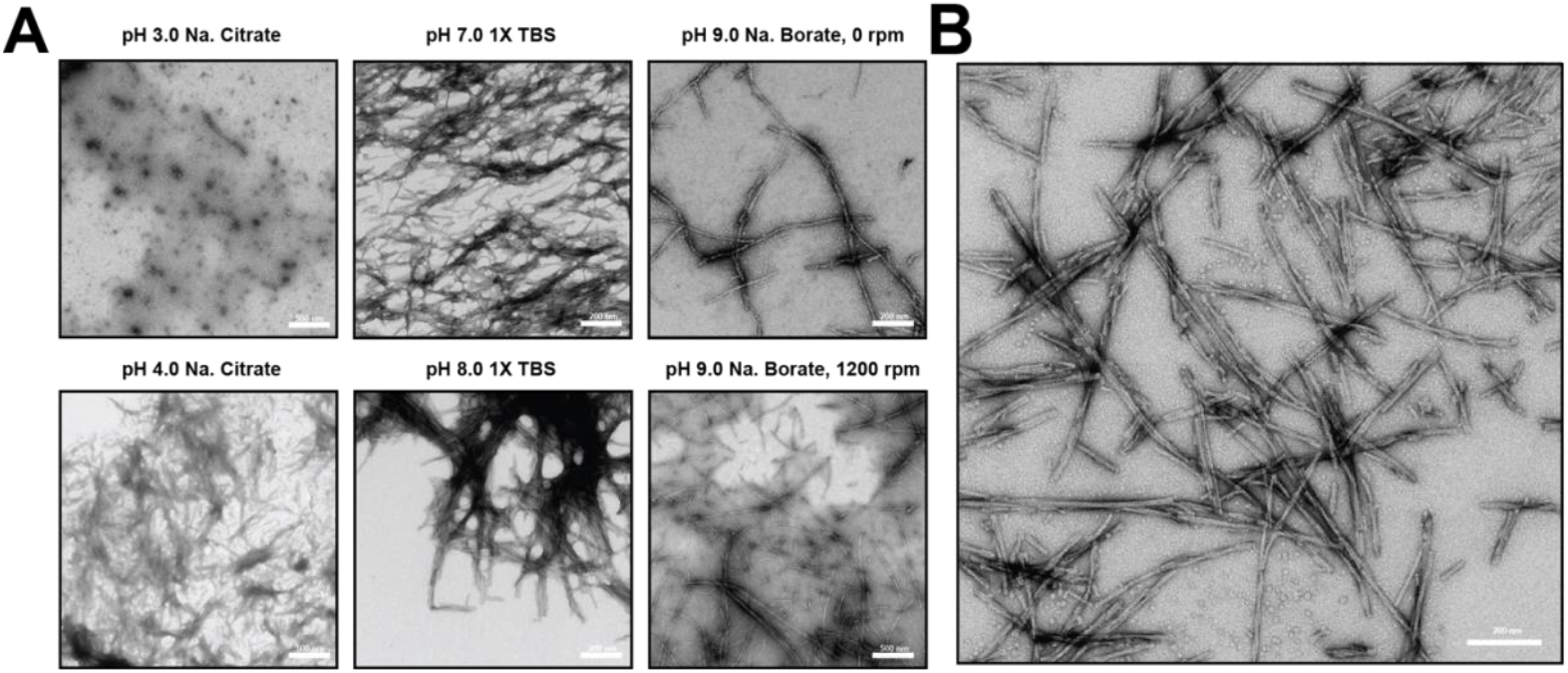
Negative stain EM images of E83Q α-Syn. A) Negative stain EM images of E83Q α-Syn from different screened conditions but not suitable for cryo-EM studies. B) Negative stain EM images of E83Q α-Syn fibrils used for cryo-EM studies.

### Cryo-EM structure of E83Q mutation

Cryo-EM micrographs of vitrified fibrils revealed a heterogeneous population of fibrils, including single-stranded, double-stranded twisted, and double-stranded ribbon-like morphologies (**Fig. 2A**). The absence of helical symmetry in the non-twisted polymorphs prevented us from determining their 3D structures using cryo-EM and helical image processing. Helical reconstruction of the twisted polymorph resulted in a 3D density map at 3.4 Å resolution (**Supplementary Table 1**). The fibrils were composed of two protofilaments arranged in C121 symmetry, and they exhibited a helical twist of 179.4 degrees and helical rise of 2.45 Å (**Fig. 2B)**. The density map was sufficient to unambiguously build an atomic model of E83Q α-Syn fibrils with well resolved sidechains for residues 36–99 (**Fig. 3A**).

**Figure 2:**
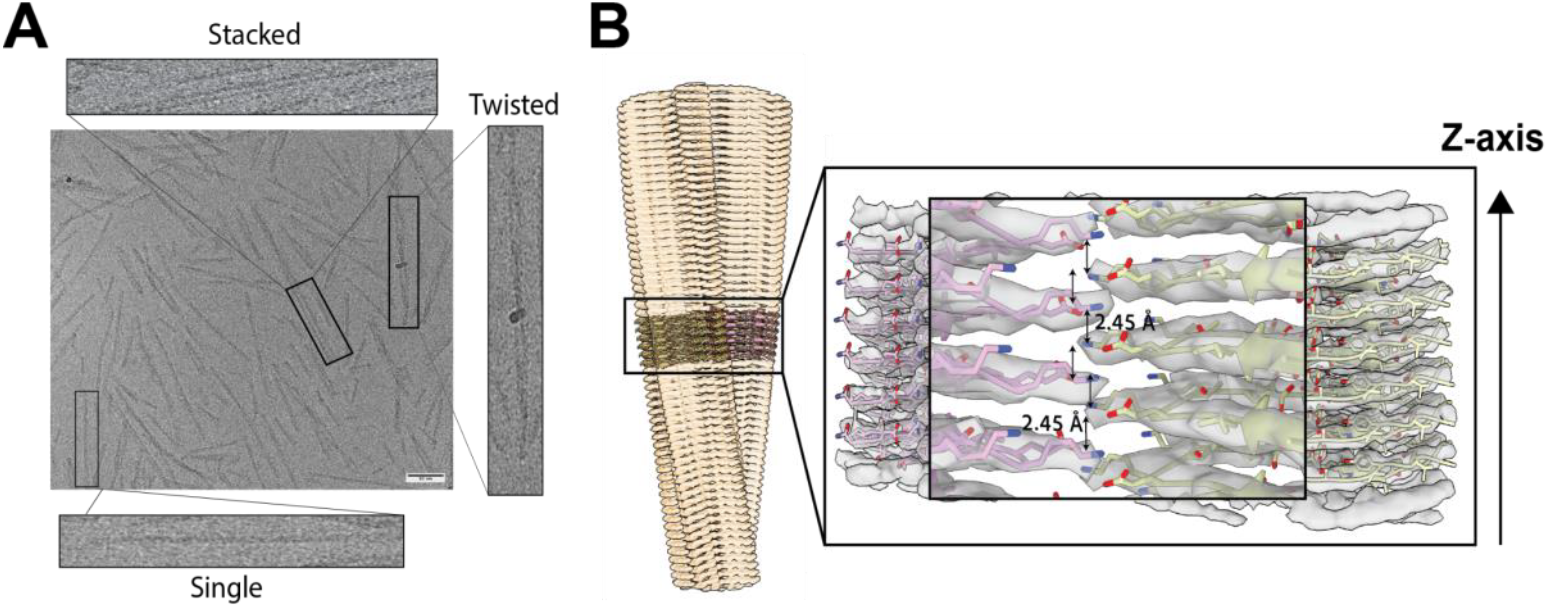
The structure of E83Q α-Syn fibrils. A) Cryo-EM image showing three different types of fibrils in the magnified boxes (stacked, straight, twisted). B) Side view of the E83Q α-Syn protofibrils, formed by two α-Syn protofilaments comprising residues 36–99, with a half-step rise of 2.45 Å.

**Figure 3:**
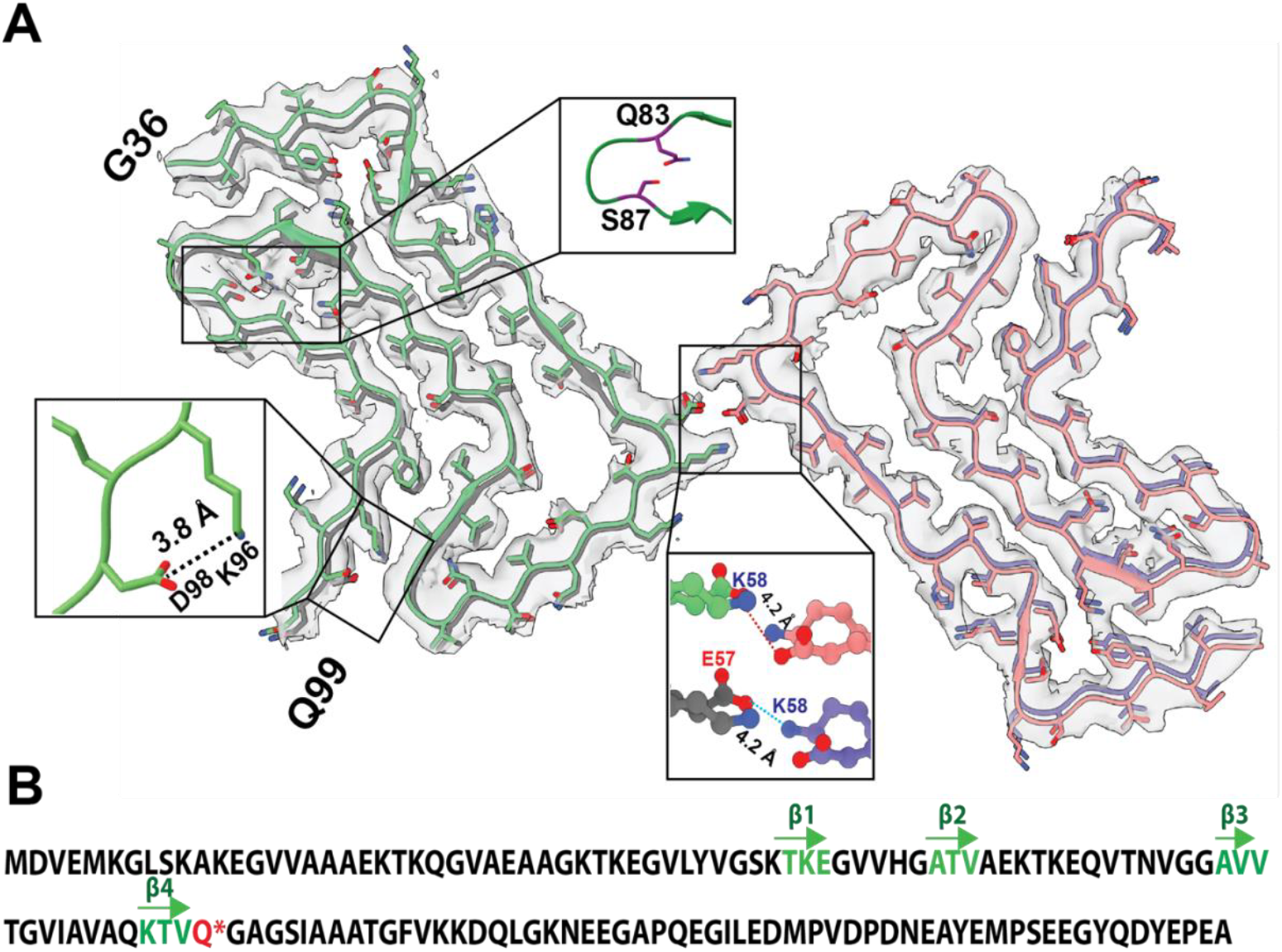
Top view of the secondary structure of E83Q α-Syn fibrils. A) Model of E83Q α-Syn fibril from residues 36 to 99, fitted into the cryo-EM density map. Salt-bridge interactions are shown between GLU57 of one protofilament and LYS58 of the other, as well as between LYS96 and ASP98 within the same protofilament. An enlargement of the mutation site (Q83) and residue S87 shows the inward orientation of residue 83. B) The β-sheets of the E83Q mutation are indicated as arrows in the α-Syn sequence.

The fibrils maintain a conserved Greek-key-like fold, with four β-strands arranged in a three-layered structure (**Fig. 3B**). The protofilament interface is stabilized by an inter-subunit salt bridge between E57 and K58, and an additional intra-filament salt bridge between K96 and D98 in the C-terminal region (**Fig. 2B, 3A**).

### Unique structural features of the E83Q α-Syn fibril and comparison to other α-Syn fibril structures

The E83Q mutation induces a unique structural feature in the fibril core. In contrast to other recombinant mutant (**Fig. 4A**), WT (**Fig. 4B**) and human brain-derived fibrils (**Fig. 4C**), the sidechain of residue 83 faces inward, away from the fibril surface, making this residue inaccessible to solvent (**Fig. 3, 4**). Interestingly, S87 also faces inward rendering it inaccessible to the solvent, as also found for A53E (PDB code 7uak) and H50Q (PDB code 6pes) mutations. The inward orientation of E83Q results in the repositioning of adjacent residues in the hydrophobic region, bringing residues 81-85 closer to the N-terminus, and blocking the N-terminal pocket **(Fig. 4**). Consequently, residues such as V37 and Y39, which are solvent-accessible in WT and other mutant fibrils, become buried in the E83Q structure.

**Figure 4:**
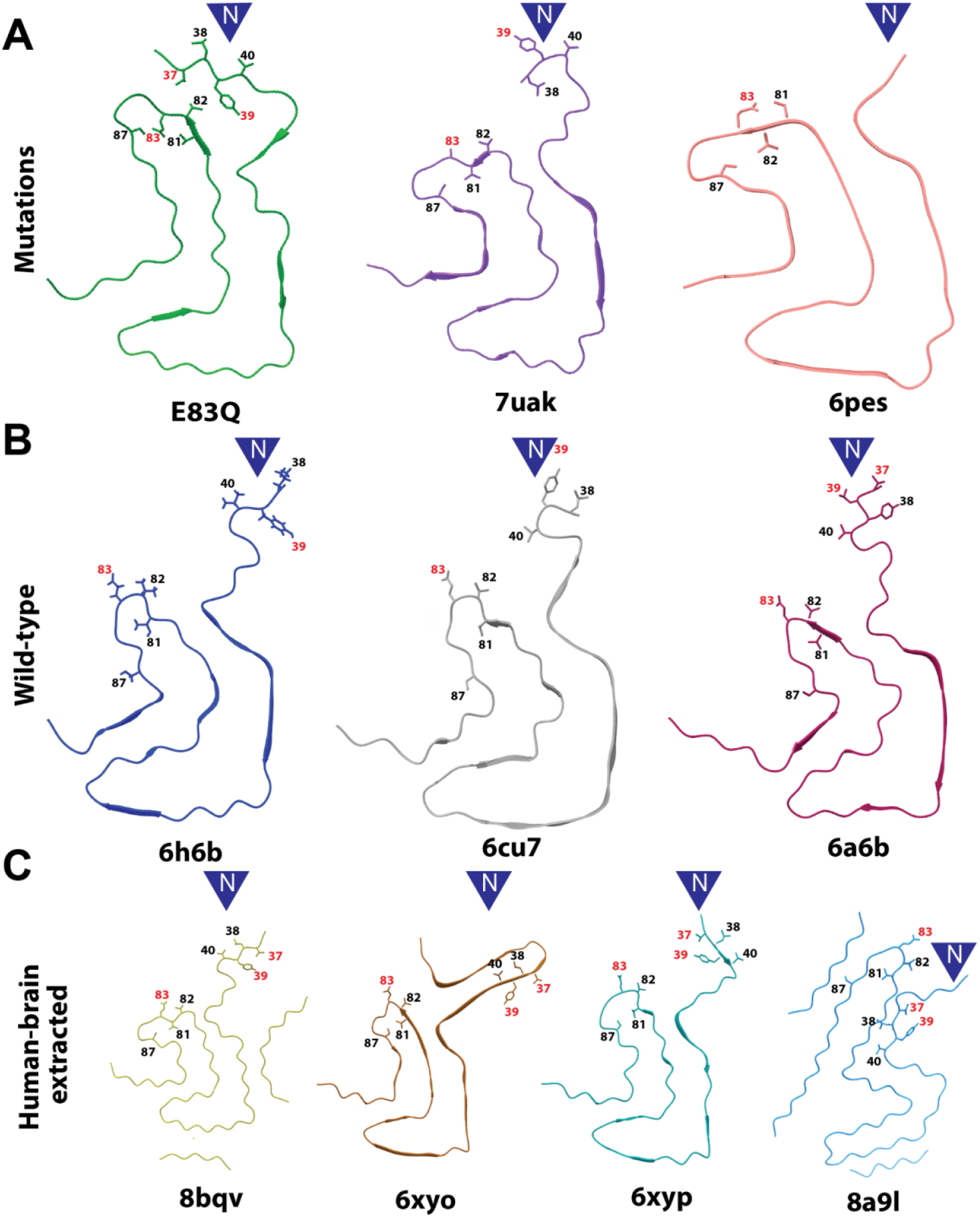
Conformation of one protofilament of different α-Syn fibrils. A) Model of α-Syn mutated fibrils for E83Q (this work), A53E (PDB code 7uak^18^) and H50Q (PDB code 6pes^19^). B) Models of wild-type α-Syn fibrils containing the Greek-key-like motif, including 1-121-truncated (PDB code 6h6b^20^), full-length (PDB code 6cu7^21^) and Full-length, N-terminal acetylated (PDB code 6a6b^22^). C) Models of human brain-extracted α-Syn fibrils from juvenile-onset synucleinopathy (JOS) (PDB code 8bqv)^23^, MSA type I (PDB code 6xyo^24^), MSA type II (PDB code 6xyp^24^) and Lewy fold from PD and DLB (PDB code 8a9l^25^). PDB codes are indicated under each model. The N-terminal pockets of the models are indicated by blue arrows.

## Discussion

Familial mutations in α-Syn act as important case studies for understanding, how subtle changes in primary sequence influence fibril structure, and ultimately, disease phenotype^16,26,27^. Many α-Syn fibrils, whether derived from *in vitro* assembly^18,19,20,21,22^or human brain tissue^23,24^, share a conserved Greek-key-like fold. Increasing evidence suggests that local structural rearrangements within this framework can significantly affect aggregation behavior and pathogenic properties. Our structural analysis of the E83Q mutation adds to this emerging view by revealing how a single charge-neutralizing mutation within the hydrophobic domain reshapes the local fibril environment without disrupting the global fold.

A defining feature of the E83Q fibril is the inward reorientation of residue 83, which contrasts with the solvent-exposed configuration observed in wild-type, other familial mutants, and brain-derived fibrils of α-Syn. This change alters the electrostatic landscape of the hydrophobic region and modifies its spatial relationship with the N-terminal domain, changing solvent mediated interactions and becoming more compact and internally stabilized. Such reorganization provides a structural rationale for the enhanced aggregation kinetics and seeding efficiency previously reported for E83Q α-Syn^16^, as reduced solvent exposure and increased local packing are likely to favor fibril nucleation and stability. This provides a structural explanation for the altered aggregation behavior observed in cellular models of E83Q α-Syn.

The altered accessibility of functionally relevant residues further distinguishes the E83Q fibril from other polymorphs. Residue 83 has been implicated in dopamine binding, an interaction known to inhibit α-Syn fibril formation ^28–30^. Therefore, burial of this site in the E83Q structure may diminish dopamine-mediated modulation of aggregation, offering a potential explanation for the more aggressive fibril behaviour observed in cell models^16^. In addition, the inward orientation of residue Ser87, reduces the accessibility of this regulatory element^31,32^ and may promote increased stability and reduced cellular turnover of E83Q fibrils. Together, these features suggest that the E83Q mutation stabilizes a fibril architecture in which key regulatory interfaces are less accessible, potentially reducing the ability of cellular factors to modulate fibril growth and turnover compared to WT or other mutant α-Syn fibrils. This altered accessibility may contribute to the distinct pathological patterns observed in E83Q-linked disease.

We here present the structure of a twisted, double-stranded α-Syn fibril polymorph. However, we also observed other classes of double-stranded, ribbon-like fibril polymorphs, and also a single-stranded form, which were all not amenable to high-resolution reconstruction. The presence of multiple fibril morphologies within the same preparation shows the pronounced structural heterogeneity of E83Q α-Syn assemblies under the conditions examined and highlights the possibility that multiple fibril conformers may co-exist *in vivo*, potentially contributing to regional or temporal variability in synucleinopathy pathology.

The structure presented here was derived from *in vitro*-assembled fibrils. It remains unclear, to which extent this architecture reflects disease-relevant assemblies in human brain. Nevertheless, in vitro– derived fibril structures have been informative for defining the range of fibril conformations accessible to α-Syn and for guiding interpretation of patient-derived structures. Resolving the cryo-EM structure of E83Q fibrils directly from patient brain tissue remains essential to establish the pathological relevance of the *in vitro* structure described here, and to define, which conformers are most closely associated with disease. Such studies may also help to explain the atypical regional distribution of pathology reported in E83Q-associated dementia with Lewy bodies, including the relatively mild involvement of the SN^15^.

In summary, our findings demonstrate that the E83Q mutation induces a distinct structural rearrangement within the conserved α-Syn fibril Greek-key framework, changing residue accessibility and domain organization in ways that are likely to influence aggregation and pathogenicity. These results reinforce the concept that disease-linked α-Syn mutations can encode specific fibril architectures, providing a structural basis for phenotypic diversity among synucleinopathies. By providing a structural basis for previously observed biochemical and cellular phenotypes, this work lays the foundation for future *in vivo* studies aimed at validating the mechanisms suggested here, and how mutation-specific architectures contribute to the heterogeneity in synucleinopathies.

## Experimental procedures

### Recombinant overexpression and purification of human E83Q α-Syn

As previously described^17^, human E83Q α-Syn was overexpressed and purified using a recombinant *E. coli* expression system. In brief, BL21(DE3) *E. coli* cells were transformed with the pT7-7 plasmid encoding E83Q α-Syn on an ampicillin-containing agar plate and incubated overnight at 37°C. The following day, a single colony was inoculated into 200 ml of Luria broth (LB) supplemented with ampicillin (100 μg/ml; AppliChem, A0839) and grown overnight at 180 rpm at 37°C. This overnight culture was then used to inoculate 6 liters of LB medium containing ampicillin (100 μg/ml), adjusting the initial absorbance at 600 nm to between 0.05 and 0.1. When the culture reached an absorbance of 0.4–0.6 at 600 nm, α-Syn expression was induced by adding 1 mM 1-thio-β-D-galactopyranoside (AppliChem, A1008), followed by an additional 4–5 hours of incubation at 180 rpm at 37°C. Cells were collected by centrifugation using a Thermo Scientific™ Sorvall LYNX 6000 Superspeed Centrifuge at 4000 rpm with a Fiberlite™ F9-6 × 1000 LEX rotor for 20 min at 4°C, and the resulting pellets were stored at −20°C. For lysis, the pellets were resuspended in 100 ml of buffer A (40 mM Tris-HCl, pH 7.5) supplemented with protease inhibitors (1 mM EDTA (Sigma-Aldrich, 11873580001) and 1 mM PMSF (AppliChem, A0999)) and subjected to ultrasonication (Vibra-Cell VCX 130, Sonics, Newtown, CT) for 8 min with 30-s on/off cycles at 70% amplitude. The lysate was centrifuged at 10,000 rpm for 30 min at 4°C, and the resulting supernatant was boiled at ∼100°C for 12–15 min. After boiling, the sample was centrifuged again at 12,000 rpm for 30 min at 4°C. The clarified supernatant was filtered through a 0.45-μm membrane and loaded into a sample loop connected to a HiPrep Q Fast Flow 16/10 column (Sigma-Aldrich, GE28-936-543). The sample was applied at 2 ml/min and eluted with buffer B (40 mM Tris-HCl, 1 M NaCl, pH 7.5) using a 0–70% gradient at 3 ml/min. Fractions were analyzed by SDS-PAGE, and those containing pure α-Syn were pooled, flash-frozen, and stored at −20°C. The pooled fractions were further purified by reverse-phase HPLC using a C4 column (PROTO 300 C4 10 μm, Higgins Analytical). The column was run with buffer A (0.1% TFA in water) and buffer B (0.1% TFA in acetonitrile), and α-Syn was eluted using a gradient from 35% to 45% buffer B over 40 min at 15 ml/min. The identity and purity of the eluted protein were confirmed by ultraperformance liquid chromatography (UPLC) and electrospray ionization mass spectrometry (ESI-MS). Fractions containing highly pure α-Syn were combined, snap-frozen, and lyophilized. The lyophilized protein was stored at −80°C until use.

### ESI-MS

ESI-MS of E83Q α-Syn for purity analysis was performed by LC-MS on the LTQ system (Thermo Scientific, San Jose, CA). Before analysis, E83Q dissolved in the buffer was desalted online by reversed-phase chromatography on a Poroshell 300SB C3 column (1.0 × 75 mm, 5 μm; on the LTQ system; Agilent Technologies, Santa Clara, CA). Protein sample (10 μl), roughly at a concentration of 0.5 to 1 μg, was injected onto the column at a flow rate of 300 μl/min and was eluted from 5 to 95% of solvent B against solvent A, linear gradient. The solvent composition was solvent A (0.1% formic acid in ultrapure water) and solvent B (0.1% formic acid in acetonitrile). MagTran software (Amgen Inc., Thousand Oaks, CA) was used for charge state deconvolution and MS analysis.

### Ultra-performance liquid chromatography (UPLC)

Protein sample (10 μl), roughly at a concentration around 1 μg, was injected to a Waters Acquity H-Class system using a C4 column (1.7 μm, 300Å, 2.1 x 150 mm) with UV detection at 214 nm and run time of 4 min (gradient 10% to 90% acetonitrile) with 0.6 mL/min flow rate.

### Screening and preparation of E83Q α-Syn fibrils for the cryo-EM studies

Monomeric E83Q α-Syn dissolved in milli-Q water was added to different buffer stocks that varied with respect to buffer composition, salt (NaCl or MgCl_2_ or KCl), pH. Different solution conditions of E83Q α-Syn that varied with concentrations under different buffer conditions were transferred to screw cap tubes having the volume of 100 µl each. Fibril formation over time and fibril morphologies suitable for cryo-EM were analyzed using negative stain EM analysis from the incubated E83Q solution conditions. Out of 12 different conditions, E83Q α-Syn fibrils at 50 mM concentration grown for 15 hours in 1X PBS buffer, pH 7.4 under 900 rpm shaking were used for the cryo-EM structural elucidation.

### Cryo-EM data collection and image processing

All sample handling and grid preparation was performed in a BSL-2 facility. 3.5 μl of fibril solution was applied to Quantifoil holey carbon grids (2/1, 200 mesh), that were glow discharged for 30 s in air to render them hydrophilic. The grids were blotted and plunge-frozen in liquid nitrogen cooled ethane, using a Leica EM GP2 plunge freezer with the sample chamber set to 4 °C and 80% humidity. Micrographs were acquired with a Titan Krios G4 (Thermo Fisher Scientific, TFS) electron microscope, operated at 300 kV acceleration voltage. Dose-fractionated electron event recordings (movies) were collected on a TFS Falcon 4i direct electron detector, using an electron dose of 50 e^−^/Å^2^. A total of 18,373 movies were recorded at a speed of around 850 movies per hour with the TFS EPU software, using a nominal magnification of 120kx, corresponding to a calibrated pixel size of 0.658 Å at the specimen level.

For data processing, EER files were imported into RELION 4.0^33^. Frames were aligned for drift correction with RELION’s internal motion correction. Frame-averaged micrographs with and without dose weighting were generated from each movie. The estimation of the contrast transfer function (CTF) for each micrograph was performed by CTFFIND4^34^ within RELION. The 16,091 micrographs with the best CTF fitting performance were selected for further processing. From these, a total of 733,876 filaments were picked using Topaz^35^. Subsequent processing was done in RELION. Fibril particles were first extracted using a box size 512 pixels and 4x4 binned to 128-pixel width, which contained 3 asymmetric units. Particles were then subjected to two-dimensional (2D) classification, and computed class averages were used to determine the helical twist and rise parameters. Using these, a three-dimensional (3D) reconstruction was computed from the generated 2D class averages. Particles in the best classes of the previous 2D classification were re-extracted with a 512-box size and binned 2x2 to a pixel size of 1.316Å for further processing. The 6,125 best helical segments were selected and subjected to particle polishing, CTF refinement, and post-processing in RELION, yielding a map at 3.4 Å resolution according to the FSC with a 0.143 threshold. Details of the data processing are summarized in Table 1.

### Model building

Atomic models were built iteratively, using Chimerax^36^, Coot^37^ and Phenix^38^ . Briefly, the structure of E83Q α-Syn was fitted into the cryo-EM density map using Coot. After refining the model in Coot and manually resolving mismatches, the model was subjected to ‘real-space refinement’ in Phenix. Side chains were then adjusted manually to the cryo-EM map using Coot Refinement in Phenix and manual adjustment were followed afterward to obtain the final model.

## Data availability

Raw cryo-EM micrographs are available at the EMPIAR database, entry numbers XXX. The 3D maps are available in the EMDB, entry number EMD-56114. Atomic coordinates are available at the PDB database with 9TPT accession number.

## Acknowledgements

We thank Hilal Lashuel for fruitful discussions. Data were collected at the Dubochet Center for Imaging (DCI) in Lausanne. The DCI is a common initiative of the EPFL and the Universities of Lausanne and Geneva. We thank the DCI for their support in cryo-EM data collection, and the Proteomics Core Facility of the EPFL for performing the mass spectrometry analysis.

## Author contributions

HS designed the study. NS and IM performed cryo-EM data collection, reconstruction and model building. BE and NS prepared the sample for mass spectroscopy. NS performed the analysis with assistance from DS. SK and ALM prepared the α-Syn fibrils. NS wrote the manuscript with assistance from AJL. All authors contributed to interpretation of the data and approved the final version of the manuscript.

## Funding and additional information

This work was in part supported by the Swiss National Science Foundation (SNF Grants CRSII5_177195 and 310030_188548) to HS. ALM was supported by the Fondation Bru, and AJL by the Parkinson Foundation Schweiz.

## Supplementary Information

**Supplementary Figure 1:**
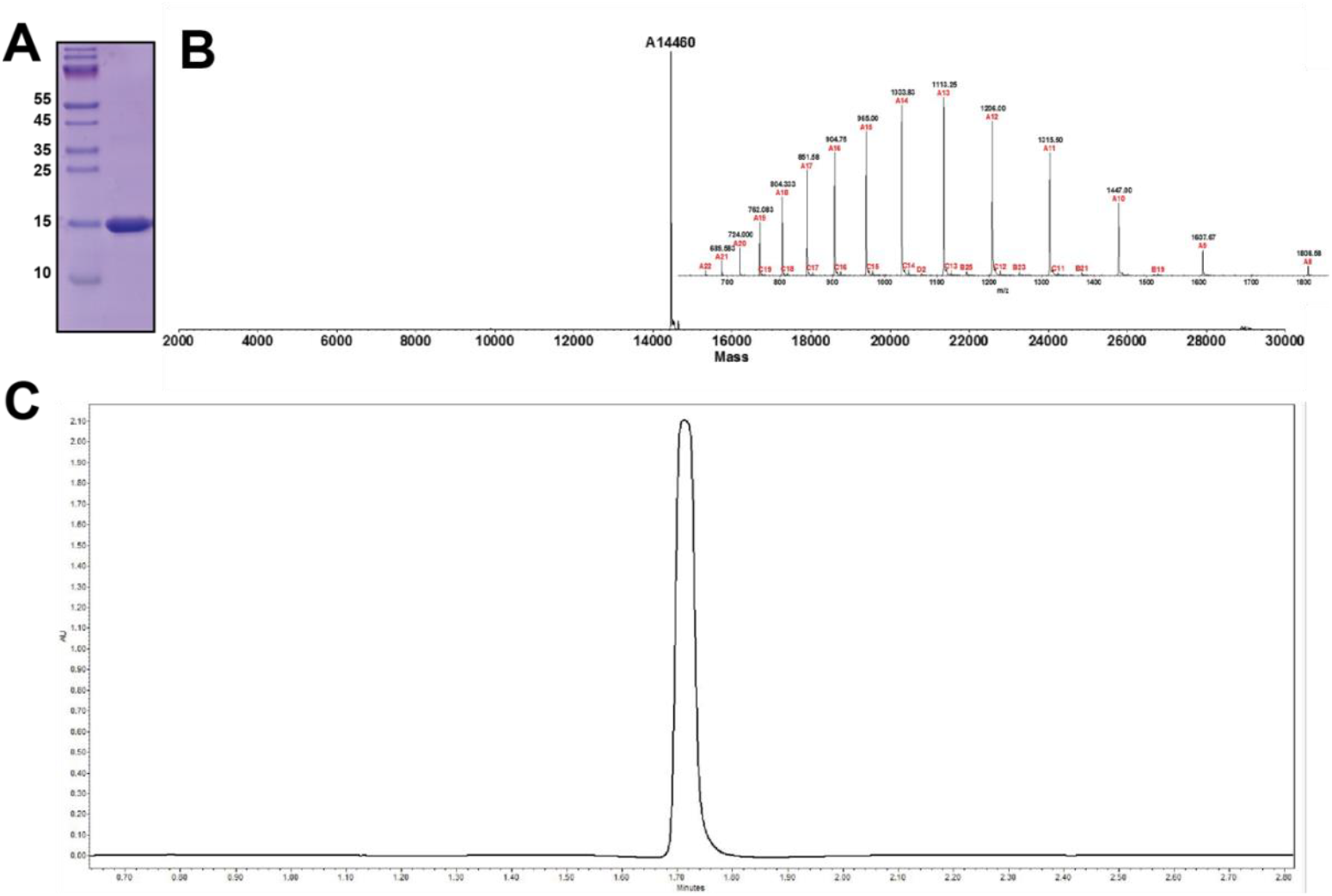
Biophysical analysis of human E83Q α-Syn fibrils. A) SDS-PAGE of human E83Q α-Syn. B) ESI-MS spectra of E83Q α-Syn. C) UPLC chromatogram of E83Q α-Syn.

**Supplementary Table 1.** Cryo-EM data collection and processing.

| **Data collection** |  |
| --- | --- |
| Microscope | TFS Titan Krios G4 |
| Acceleration Voltage [kV] | 300 |
| Electron source | cold-FEG |
| Detector | TFS Falcon 4i |
| Nominal magnification | 120 kx |
| Pixel size [Å] | 0.658 |
| Total electron dose [e<sup>-</sup>/Å<sup>2</sup>] | 50 |
| Exposure time [s] | 4.2 |
| Defocus range [μm] | -0.8 to -2.0 |
| **Reconstruction** |  |
| No. of micrographs collected | 18,373 |
| No. of filaments picked | 733,876 |
| Initially extracted segments | 1,294,708 |
| No. of segments for 3D auto-refine | 6,125 |
| Resolution [Å] | 3.43 |
| Map sharpening B-factor [Å<sup>2</sup>] | -80 |
| Symmetry for final map | C121 |
| Helical rise [Å] | 2.45 |
| Helical twist [°] | 179.43 |
| **Atomic model** |  |
| No. of non-hydrogen atoms | 5,352 |
| No. of protein residues | 36 – 99 |
| R.m.s. deviations |  |
| Bond lengths [Å] | 0.003 |
| Bond angle [°] | 0.650 |
| Ramachandran plot statistics [%] |  |
| Preferred | 89.6 |
| Allowed | 10.4 |
| Outlier | 0.00 |
| Rotamer outliers [%] | 0.00 |
| MolProbity score | 1.95 |
| Clashscore | 7.01 |

